# Lack of Cdk5 activity is involved on Dopamine Transporter expression and function: Evidences from an animal model of Attention-Deficit Hyperactivity Disorder

**DOI:** 10.1101/2021.03.19.436226

**Authors:** Fernández Guillermo, Krapacher Favio, Ferreras Soledad, Quassollo Gonzalo, Mari Macarena Mariel, Pisano María Victoria, Montemerlo Antonella, Rubianes María Dolores, Bregonzio Claudia, Carlos Arias, Paglini María Gabriela

**Author notes:** **Corresponding author:** María Gabriela Paglini, Ph.D., Laboratory of Neurophysiology, Instituto de Investigación Médica Mercedes y Martín Ferreyra, INIMEC-CONICET, Universidad Nacional de Córdoba, Friuli 2434, 5016 - Córdoba Argentina., (ext. 105).

## Abstract

Attention deficit/Hyperactivity disorder (ADHD) is one of the most diagnosed psychiatric disorders nowadays. The core symptoms of the condition include hyperactivity, impulsiveness and inattention. The main pharmacological treatment consists of psychostimulant drugs affecting Dopamine Transporter (DAT) function. We have previously shown that genetically modified mice lacking p35 protein (p35KO), which have reduced Cdk5 activity, present key hallmarks resembling those described in animal models useful for studying ADHD. The p35KO mouse displays spontaneous hyperactivity and shows a calming effect of methylphenidate or amphetamine treatment. Interestingly, dopaminergic neurotransmission is altered in these mice as they have an increased Dopamine (DA) content together with a low DA turnover. This led us to hypothesize that the lack of Cdk5 activity affects DAT expression and/or function in this animal model. In this study, we performed biochemical assays, cell-based approaches, quantitative fluorescence analysis and functional studies that allowed us to demonstrate that p35KO mice exhibit decreased DA uptake and reduced cell surface DAT expression levels in the striatum (STR). These findings are supported by *in vitro* observations in which the inhibition of Cdk5 activity in N2a cells induced a significant increase in constitutive DAT endocytosis with a concomitant increase in DAT localization to recycling endosomes. Taken together, these data provide evidences regarding the role of Cdk5/p35 in DAT expression and function, thus contributing to the knowledge of DA neurotransmission physiology and also providing therapeutic options for the treatment of DA pathologies such as ADHD.

## INTRODUCTION

Attention Deficit Hyperactivity Disorder (ADHD) is a complex, chronic and highly heritable neurodevelopmental disorder with typical onset in childhood and known persistence into adulthood (Sharma and Couture, 2014). Like other neuropsychiatric disorders, ADHD presents a spectrum of behavioral alterations with motor hyperactivity features, impulsivity, and/or inattention; offering the diagnostic criteria used for diagnosis (American Psychiatric Association, 2013). So far, the most effective pharmacotherapy for the disorder comprises long-term treatments with stimulant drugs, such as amphetamine (AMPH) and methylphenidate. These drugs enhance Dopamine (DA) neurotransmission by altering Dopamine Transporter (DAT) normal function (Faraone et al., 2005; German et al., 2015; Hong and Amara, 2013). DAT is a key plasma membrane-protein regulating dopaminergic transmission, both spatially and temporally, since it removes released DA from the synaptic cleft by a reuptake mechanism (German et al., 2015). Given the essential role of DAT in dopaminergic transmission, several studies support its participation in ADHD. Consistently, results obtained using genetically modified animals like DAT knockout (DAT-KO) and cocaine-insensitive DAT (DAT-CI) mice, highlight the role of the transporter in phenotypes related to ADHD (Federici et al., 2014; Hawi et al., 2015; Leo and Gainetdinov, 2013; Li et al., 2014; Napolitano et al., 2010; O’Neill and Gu, 2013; Sakrikar et al., 2012). Considering that DAT is functional only when it is expressed at the plasma membrane, its surface availability is dynamically and tightly regulated by endocytic trafficking (Bermingham and Blakely, 2016; Eriksen et al., 2010; Melikian, 2004). It has been shown that DAT endocytosis is mediated by a clathrin-dependent mechanism (Sorkina et al., 2005) and also that Cdk5 kinase plays an essential role in regulating synaptic vesicle endocytosis by phosphorylation of Dynamin I, a protein that participates in clathrin-dependent internalization (Tan et al., 2003; Tomizawa et al., 2003).

Cdk5 is a proline-directed serine/threonine kinase that is activated by its association with a regulatory subunit, either p35 or p39, both of which are brain-specific (Ishiguro et al., 1994; Lew and Wang, 1995; Tang et al., 1995; Tsai et al., 1994). Cdk5/p35 complex has been involved in several processes, including neuronal migration, axonal and dendritic formation (Paglini et al., 1998; Shah and Lahiri, 2017; Shah and Rossie, 2018), neurotransmitter release (Kim and Ryan, 2010), endocytosis (Nguyen and Bibb, 2003; Tan et al., 2003), dopaminergic signaling (Svenningsson et al., 2004) as well as in structural plasticity associated to drugs of abuse (Ferreras et al., 2017; Mlewski et al., 2008, 2016). Furthermore, Cdk5 activity is tightly regulated in neurons under physiological conditions. Dysregulation of Cdk5 activity gives rise to various neurodevelopmental, neurological and neuropsychiatric disorders, such as Alzheimer’s, Parkinson’s and Huntington’s diseases, ADHD, schizophrenia and stress-induced hippocampal memory deficits, among others (Ikiz and Przedborski, 2008; Kawauchi, 2014; Piccini et al., 2015; Rei et al., 2015; Shah and Lahiri, 2014). In line with these observations, results from our laboratory showed that mice lacking a proper Cdk5 activity recapitulate some key features present in ADHD. By using a mutant mouse lacking p35 (p35KO) (Chae et al., 1997), which displays reduced Cdk5 activity; we demonstrated that juvenile mice mimic key hallmarks of the human disease, including spontaneous hyperactivity and the decrease of locomotor activity after AMPH or methylphenidate treatment; behavioral response defined as “*paradoxical response to psychostimulants*”. Moreover, p35KO mice also exhibited increased DA synthesis and content and decreased DA degradation, resulting in a low DA turnover in basal conditions (Krapacher et al., 2010). These findings together with those previously reported (Drerup et al., 2010) support the validity of the p35KO mouse as an animal model useful for the study of ADHD (de la Peña et al., 2017). Although Cdk5 activity plays a critical role in dopaminergic transmission, dysregulation of Cdk5 and its effects on DAT expression, trafficking and function remain largely unexplored.

In the present study we tested the hypothesis that the lack of p35-activated Cdk5 activity affects DAT expression and/or function in p35KO mice. We demonstrate that dopaminergic activity is altered in the brain of this animal model. Decreased DA uptake and reduced cell surface DAT expression levels were observed in the STR of these mutant animals. These findings are supported by *in vitro* observations, in which the lack of Cdk5 activity induced a significant increase in constitutive DAT endocytosis. Given the central role of DAT in dopaminergic signaling, identifying molecular mechanisms underlying DAT membrane expression, trafficking and function not only contributes to broaden our understanding of DA neurotransmission but also provides therapeutic options for the treatment of DA pathologies such as ADHD.

## MATERIALS AND METHODS

### Animals

Wild type (WT) and p35 knockout (p35KO) mice were generated by breeding heterozygous mutants (kind gift of Dr. L.H. Tsai) (Chae et al., 1997), maintained in a C57BL/6J background via brother-sister mating in the vivarium of the INIMEC-CONICET-UNC (Cordoba, Argentina). Mice were weaned at 21 postnatal days and housed up to six per cage under 12 h light /12 h dark cycle, controlled temperature (22°C), and free access to food and water. All experiments were performed using male animals at the age of 24-26 days. All animal procedures and care were approved by the National Department of Animal Care and Health (SENASA–ARGENTINA) and were in compliance with the National Institute of Health general guidelines for the Care and Use of Laboratory Animals. All experimental protocols were reviewed and approved by the Institutional Animal Care and Use Committee (CICUAL) at INIMEC-CONICET-UNC. Efforts were made to minimize animal suffering and to reduce the number of animals used.

### Antibodies and Drugs

Sources of primary antibodies: Hemaglutinin (HA), Y11 Santa Cruz Biotechnology (Santa Cruz, CA, USA) (1:400), LAMP-1, H4A3, Iowa Hybridomic bank (Iowa, IA, USA) (1:500), DAT, AB15344 Millipore (Billerica, MA, USA) (1:800, recognizes two bands at 55 and 75 kDa in rat brain tissue and one band at 55 kDa in mouse brain tissue); α-tubulin, DM1A Sigma-Aldrich (St. Louis, MO, USA) (1:5000, recognizes α-subunit of tubulin at 50 kDa); Na^+^K^+^-ATPase, 464.6(6H) GeneTex (Irvine, CA, USA) (1:50000, recognizes Sodium/Potassium ATPase Alpha-1 at 100 kDa); glyceraldehyde 3-phosphate dehydrogenase (GAPDH) AM4300 Invitrogen (San Diego, CA, USA) (1:5000, recognizes a 37 kDa band corresponding to GAPDH in different cell lines and tissues tested). Sources of secondary antibodies used in western blot: goat anti-mouse IgG IRDye 680/ goat anti-rabbit IgG IRDye 800, Li-Cor Biosciences (Lincoln, NE, USA) (1:10000). Sources of secondary antibodies used in inmunocytochemistry: goat anti-rabbit IgG Alexa Fluor 488 (GAR-488) and goat anti-rabbit IgG Alexa Fluor 568 (GAR-568), ThermoFisher Scientific (Waltham, MA, USA) (see working dilutions in sections below).

D-Amphetamine (AMPH) (Parafarm, Buenos Aires, Argentina) and roscovitine (ROSCO) (Calbiochem, San Diego, CA, USA) were used in this study.

### Cell line and DNA constructs

Mouse Neuro 2a (N2a) cells (CCL-131), acquired from American Type Culture Collection (Manassas, VA) were cultured at 37°C, 5% CO_2_ in Dulbecco’s Modified Eagle Medium (DMEM, ThermoFisher Scientific, Waltham, MA, USA) supplemented with fetal bovine serum (FBS) 5% (NATOCOR, Córdoba, Argentina), glutamine 2 mM, penicillin 10,000 U/ml, and streptomycin 10 mg/ml (ThermoFisher Scientific, Waltham, MA, USA). N2a cells were transiently transfected using Lipofectamine 2000 (ThermoFisher Scientific, Waltham, MA, USA) according to the manufacturer’s recommendations and as described (Ferreras et al., 2017). Briefly, one hour before transfection, N2a cells grown on coverslips were transferred to 35 mm culture plates containing 1 ml of serum and antibiotics free Opti-MEM (Gibco, ThermoFisher Scientific, Waltham, MA, USA). Then, DNA constructs were gently mixed in a tube containing 125 µl of Opti-MEM. Simultaneously, 6 µl of Lipofectamine 2000 were gently mixed with 125 µl of Opti-MEM. After 5 min. of incubation, the diluted DNA was combined with the diluted Lipofectamine 2000 and the mix was incubated for 20 min. at room temperature. Then, the mixture was added to the N2a cells in the 35 mm culture plates and incubated for 2 hs. at 37°C. After that, the medium was removed and replaced by serum-free DMEM. Experiments were performed 24 hs. after transfection. For antibody feeding method (AFM) experiments, N2a cells were transfected with a DAT containing a HA epitope (YPYDVPDASL) in the second extracellular loop (DAT-HA) (Hong and Amara, 2013). This construct was a kind gift from Dr. Susan Amara (National Institute of Mental Health, Bethesda, MD, USA). In colocalization experiments, early and recycling endosomes markers, Rab5-GFP and Rab11-GFP, respectively, were used. These plasmids were a kind gift from Dr. José Luis Daniotti (CIQUIBIC-Universidad Nacional de Córdoba, Córdoba, Argentina).

### Surface DAT biotinylation assay

For the assessment of surface DAT levels in synaptosomes from striatal tissue, crude synaptosomal fractions were obtained as described (Mlewski et al., 2008). Briefly, naïve p35KO and WT mice were sacrificed by cervical dislocation and the STRs were quickly dissected out on ice-cooled dishes and homogenized in ice-cold Sucrose Buffer (0.32 M sucrose, 10 mM HEPES and 1 mM EDTA pH 7.4, 1 µg/ml aprotinin, 1 µg/ml leupeptin, 100 µg/ml PMSF, 1 µg/ml pepstatin, and 0.2 mM sodium orthovanadate; all chemicals were from Sigma-Aldrich, St. Louis, MO, USA) using a small Potter glass-Teflon homogenizer. The resulting homogenate was passed three times trough a 30G syringe. The sample was then centrifuged twice at 1000 G for 5 min at 4 °C. The supernatant (total homogenate, HT) was centrifuged at 12500 G for 20 min at 4°C. The resulting pellet (crude synaptosomal fraction) was resuspended in 1 ml of PBS/Ca^+^/Mg^+^ buffer and protein content was quantified by DC protein assay kit (Bio-Rad Laboratory, Hercules, CA, USA). 500 µg of synaptosomes were incubated 60 min at 4 °C with sulfo-NHS-biotin (1.5 mg/ml) in 500 µl of PBS/Ca^+^/Mg^+^ pH 7.4 (ThermoFisher Scientific, Waltham, MA, USA). Biotinylation reaction was quenched by incubating the synaptosomes with 1 vol of PBS containing glycine 100 mM for 15 min. Subsequently, the sample was washed twice with the same buffer. Synaptosomes were then resuspended in lysate buffer (1% Triton X-100, 1% Nonidet P-40, 10% glycerol in TBS buffer pH 7.4 supplemented with 1 µg/ml aprotinin, 1 µg/ml leupeptin, 100 µg/ml PMSF, 1 µg/ml pepstatin and 0.2 mM sodium orthovanadate; all chemicals were from Sigma-Aldrich, St. Louis, MO, USA). After that, samples were sonicated twice (10 sec x 30 mA). An aliquot of this fraction was separated (synaptosomal total DAT). Samples (300 µg) were then incubated with streptavidin beads 100 µl (ThermoFisher Scientific, Waltham, MA, USA) overnight at 4° C. Beads were washed five times with lysate buffer and biotinylated proteins were eluted with Laemmli buffer (62,5 mM Tris-HCl, pH 6.8, 20% glycerol, 2% SDS, and 5% β-mercaptoethanol) for 30 min at RT.

### Surface DAT biotynilation after AMPH treatment

Striatal synaptosomes (500 µg) from p35KO and WT mice, following the same procedure as describe above, were resuspended in Krebs Ringer Buffer (KRB) containing 145 mM NaCl, 2.7 mM KCl, 1.2 mM KH_2_PO_4_, 1.2 mM CaCl_2_, 1 mM MgCl_2_, 10 mM glucose, 0.255 mM ascorbic acid, and 24.9 mM NaHCO_3_. Samples were then incubated with AMPH (10 µM) or VEH for 30 min at 37°C in gentle agitation. The reaction was then stopped with two ice-cold KRB washes. Subsequently, the biotinylation procedure was carried out as described in the previous section.

### Western Blot

Protein immunoblotting was performed as previously described (Ferreras et al., 2017). HT (25 µg), synaptosomal total DAT (25 µg) and half of each biotinylated fraction were separated into 10% SDS-PAGE gels electrophoresis and transferred to nitrocellulose membranes (Bio-Rad Laboratory, Hercules, CA, USA). Subsequently, membranes were first incubated 60 min at RT with blocking buffer (5% skim milk in TBS tween 0,05%) and then incubated over night at 4 °C in agitation with the following antibodies: anti-DAT, anti-tubulin, anti-Na-K-ATPase and anti-GAPDH. Membranes were then washed and incubated 60 min at RT with the appropriate Licor IRDye secondary antibody. Protein signals were detected using an Odissey scanner (LI-COR Biosciences, Lincoln, NE, USA). Blot-band densitometries were quantified using the default protocol of gels plug-in of FIJI software (https://imagej.nih.gov/ij/docs/menus/analyze.html#gels). Relative protein expression was calculated as the relation of the densitometry of the protein of interest and its corresponding control. Results are shown as a percentage of control treatments.

### Dopamine uptake in p35KO and WT mice STRs: amperometric experiments

Naïve p35KO and WT mice were sacrificed by cervical dislocation and the STRs were quickly dissected on ice, chopped with a razor blade and homogenized in 1.1 ml of cold KRB saturated with a gas mixture of 95% O_2_ and 5% CO_2._ Subsequently, DA uptake was carried out as described below.

#### Electrochemical procedure

DA uptake was evaluated by amperometric measurements with a glassy carbon electrode (GCE) modified with a dispersion of multi walled carbon nanotubes (Nano-Lab, Waltham, MA, USA) functionalized with polietilenimine (Sigma, St. Louis, MO, USA) (CNT-PEI) as sensing platform. The electrochemical measurements were performed with TEQ-04 potentiostats. The electrodes were inserted into the cell (BAS, Model MF-1084) through holes in its Teflon cover. A platinum and an Ag/AgCl wire in 3M KCl were used as counter and reference electrodes, respectively. All potentials measured by the working electrode (BAS, Model RE-5B) were referred to the Ag/AgCl electrode. A magnetic stirrer provided the convective transport during the amperometric measurements.

The experiments were carried out in a KRB solution saturated with a gas mixture (95% O_2_ and 5% CO_2_) at 37°C, by applying +0.250 V and monitoring the current response in the absence or in the presence of 500 µl of homogenized striatal samples. The oxidation current signal value was measured initially (*i*_*0*_) and after 300 seconds (*i*_*300*_) of the addition of 5.0 × 10^−7^ M DA (Sigma, St. Louis, MO, USA). The decrease in the oxidation current signal was used to calculate DA pmols taken up by DAT. DA reuptake was finally expressed as pmol of DA/µg of total protein/min. In order to perform the measurements as quickly as possible, the total protein content was measured after the experiment.

The glassy carbon electrode modified with carbon nanotubes dispersed in polyethylenimine (GCE/CNT-PEI) was prepared as described by Rubianes and Rivas (Rubianes and Rivas, 2007). GCE was polished with alumina slurries of 1.00, 0.30 and 0.05 µm for 1 min each and sonicated in water for 30 s, and then modified by drop coating with an aliquot of 20 µL of CNT-PEI and allowed to dry for 90 min. The dispersion of CNT-PEI was obtained by sonicating 1.0 mg of MWCNTs within 1.0 mL of 1.0 mg/mL PEI solution (prepared in 50:50 v/v ethanol/water) for 5.0 min with an ultrasonic probe.

### Antibody feeding method (AFM) and pharmacological treatments

In order to perform AFM, a quantitative immunofluorescence technique (Hong and Amara, 2013; Sorkina et al., 2006), N2a cells were grown in coverslips and transfected with DAT-HA. 24 hs after transfection, cells were incubated with anti-HA (Y11) 20 µg/ml and ROSCO 10 µM or DMSO (vehicle) in FBS-free DMEM for 30 min at 18 °C. After that, DMSO treated cells were incubated with AMPH 10 µM (AMPH group) or VEH (Control group), meanwhile, ROSCO treated cells were maintained with this drug (ROSCO group) for 60 min at 37 °C. Subsequently, cells were washed with PBS and fixed for 20 min with 4% paraformaldehyde (Riedel-de Haën, Sigma-Aldrich Laborchemikalien GmbH, Seelze, Germany) in 100 mM Phosphate Buffered Saline (PBS) at room temperature (RT). In order to label surface DAT-HA, cells were incubated with a secondary anti-rabbit antibody Alexa Fluor-568 (80 µg/ml) in PBS containing Bovine Serum Albumin (BSA) 1% at RT for 60 min. Cells were then washed with PBS and permeabilized with Triton X-100 0.2% BSA 1% in PBS at RT. Cells were then incubated with a secondary anti-rabbit antibody Alexa Fluor-488 (2 µg/ml) in PBS containing BSA 1% at RT for 60 min to label endocyted DAT-HA. After incubation, cells were washed and mounted on slides using FluorSave (Calbiochem Merck KgaA, Darmstadt, Germany). Subsequently, high-resolution images of N2a cells were acquired in FV-1000 laser-scanning confocal microscope (Olympus, Tokyo, Japan) in two channels (excitation wavelength: 494/578, emission wavelength: 517/603) using an oil immersion 63X (NA 1.4) objective (PlanApo, Olympus). Optical sections were acquired at 0.5 µm intervals from the top to bottom of cells and data was imported into ImageJ (U.S. National Institutes of Health, Bethesda, MD, USA) for analysis. For each cell, an average of integrated density value (product of Area and Mean Gray value) from three medial Z-stacks was obtained both for intracellular DAT signal as well as superficial DAT. Background intensity was subtracted from images prior to quantification. After that, the ratio of intracellular to surface DAT signal was calculated as an internalization index for statistical analysis.

### Colocalization experiments and pharmacological treatments

N2a cells were grown in coverslips and co-transfected with plasmids encoding DAT-HA and Rab5-GFP or Rab11-GFP. After 24 hs of expression, cells were incubated with ROSCO (10 µM) or AMPH (10 µM) for 30 min in FBS free DMEM. Cultures were immediately washed with PBS and fixed for 20 min with 4% paraformaldehyde in 100 mM PBS at RT. Subsequently, cells were permeabilized with Triton X-100 0.2 mM in PBS. After three PBS washes, cells were incubated with BSA 5% in PBS for 60 min and then incubated 60 min at RT with primary antibody anti-HA (Y11) to label DAT-HA. Cells were then incubated with anti-rabbit Alexa-Fluor 568 (1:400) for 60 min, washed three times and mounted onto slides using FluorSave.

In DAT/LAMP-1 colocalization experiments, cells were transfected with DAT-HA. After 24 hs of expression cells were incubated with ROSCO (10 µM) for 30 min and then processed as described above. Endogenous LAMP-1 was labeled with a primary antibody anti-LAMP-1 and a secondary antibody AlexaFluo-488 (1:400).

Subsequently, images were acquired in FV-300 laser-scanning confocal microscope (Olympus, Tokyo, Japan) in two channels (excitation wavelength: 494/578, emission wavelength: 517/603) using an oil immersion 63X (NA 1.4) objective (PlanApo, Olympus). In these experiments, optical sections were recorded in 0.25 µm intervals and data were imported to ImageJ for quantitative colocalization analysis. Given that serum deprivation in N2a cells promotes the development of lamellar structures similar to neurites (López-Maderuelo et al., 2001), fluorescence intensity of each endosomal marker protein was quantified only in neurite-like processes. In view of the low number of LAMP-1 positive vesicles in neurite-like processes, for LAMP-1/DAT colocalization analysis, quantification was done using the whole cell.

Images were deconvoluted using PSF Generator and Deconvolution lab plug-in (Biomedical Imaging group) (Sage et al., 2017). For each cell, a focal plane with well-defined endomembrane structures was selected for analysis and colocalization was quantified in the 15 µm distal portion of neurites with the ImageJ plugin JaCoP (Bolte and Cordelières, 2006). Threshold for DAT and endomembrane marker signals were defined for each cell in order to avoid colocalization overestimation. The colocalization coefficient selected for the analysis was Manders M1 (MCC). This was defined as the summed intensities of pixels from endosomal marker images for which the intensity in DAT images was above zero compared to the total intensity in the endosomal marker images (Bolte and Cordelières, 2006).

### Statistical analysis

Statistical analyses were performed using R software (R Core Team, 2020). Assumptions of homoscedasticity and normality were tested using Levene and Shapiro-Wilks tests, respectively. In those cases in which the assumptions were met, we used unpaired student *t*-test, one-way or two-way ANOVA as appropriate followed by Tukey post-hoc test (Type I error set at 0.05). Data that did not meet these assumptions were analyzed with the non-parametric tests Kruskal-Wallis followed by Mann-Whitney’s test when necessary.

## RESULTS

### Mice lacking Cdk5-activator p35 display decreased DA uptake in the striatum

We have previously shown that DA content is dramatically augmented in the STR of mutant mice lacking p35. In spite of the high DA concentration, these mutant mice produce less DOPAC than their WT littermates, resulting in a low DA turnover in basal conditions (Krapacher et al., 2010). Considering that DA homeostasis is maintained mainly by DAT through monoamine reuptake, we analyzed DAT expression and function in the STR of these mice.

We first determined DAT expression levels in total striatal homogenates and synaptosomes from juvenile naïve WT and p35 KO mice. Western blots analyses showed no differences in DAT expression levels in total homogenates (t-test: t = 0.53972, df = 6, p = 0.6088) or synaptosomal fractions (t-test: t = 0.03702, df = 6, p = 0.9717) between WT and p35KO mice (Fig. 1a). Simultaneously, to test the functional consequences of the lack of p35-activated Cdk5 activity in DA uptake, we performed amperometry experiments in the STR of p35KO and WT mice. Homogenized STR tissues from both genotypes were incubated with DA and the uptake of DA was recorded for 5 min. The quantitative analysis showed a dramatic decrease of 50% in DA uptake in p35KO preparations compared to WT controls (Fig. 1b) (t-test: t = -3.1773, df = 6, p < 0.05).

**Figure 1:**
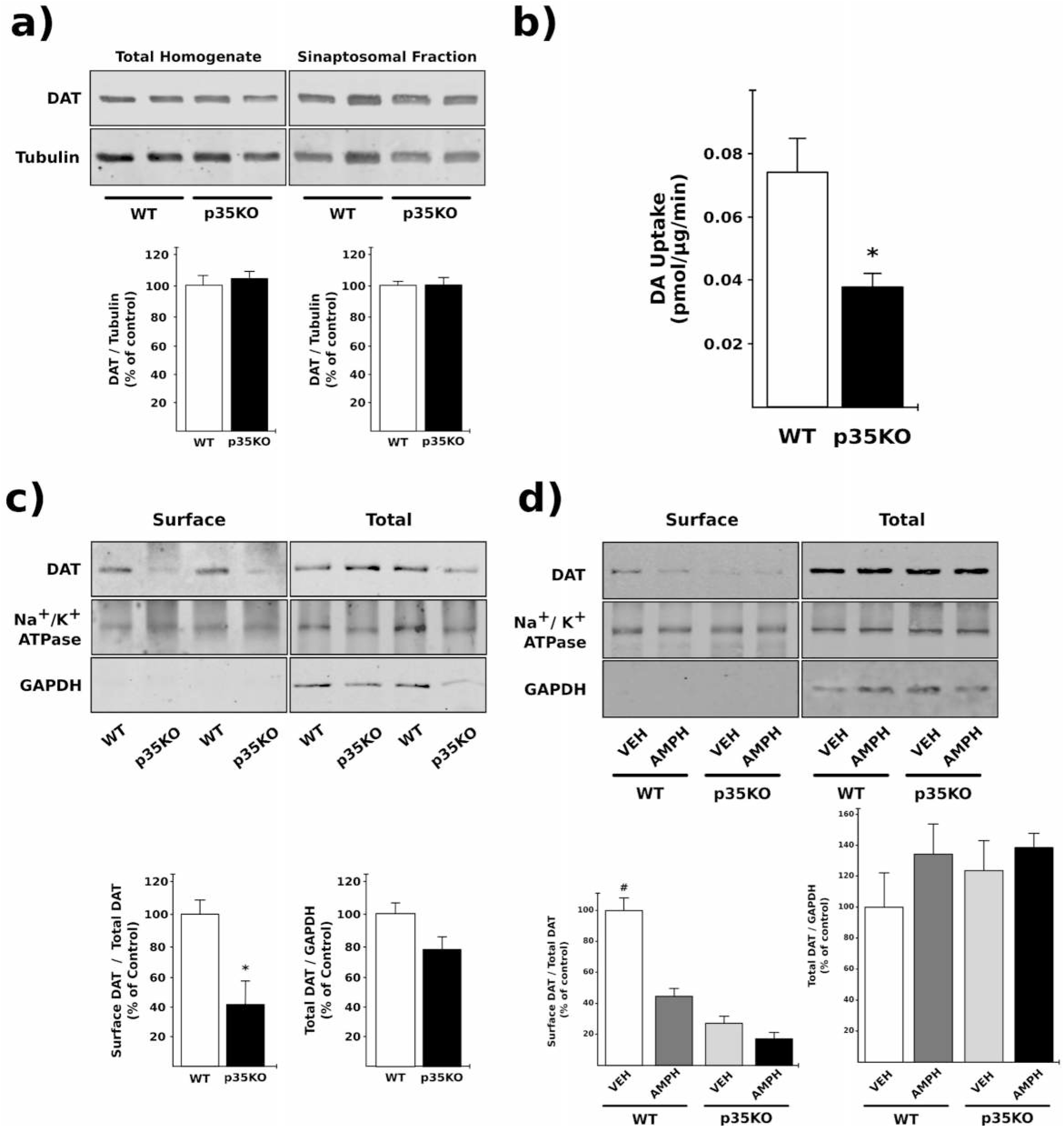
p35KO mice exhibit decreased DA uptake and DAT cell surface expression in the STR. (a) Representative western immunoblots showing DAT expression levels in total STR homogenates and synaptosomal fractions from WT and p35KO mice (measured by DAT/Tubulin ratio; mean ± SEM, Student t-test, n= 4). (b) DAT activity in STR homogenates from p35KO and WT mice. Striatal tissue was homogenized in KRB and DA uptake assay was performed by an amperometric technique using a glassy carbon electrode. Results are expressed as pmol of DA/µg of protein/min (mean ± SEM, Student t-test, * indicates p < 0.05, n= 4). (c) Surface protein biotinylation experiments in striatal synaptosomes from WT and p35KO mice. Representative western immunoblots of biotinylated (surface) and non-biotinylated (total) DAT expression levels in striatal synaptosomes from WT and p35KO mice (upper panels). Surface DAT levels were measured as surface/total DAT ratio. Note the decreased expression of DAT in the surface fraction of p35KO mice. Medial panels show surface and total Na^+^/K^+^ ATPase expression levels and bottom panels, GAPDH levels. Total DAT levels were quantified measuring total DAT/GAPDH ratio (mean ± SEM, Student t-test, *\*p* < 0.05, n= 3-4). (d) AMPH effect on DAT surface expression in striatal synaptosomes from WT and p35KO mice. Representative western blots of biotinylated (surface) and non-biotinylated (total) DAT expression in striatal synaptosomes from animals of both genotypes incubated for 30 min with AMPH or VEH. DAT surface expression was measured as surface/total ratio. Total DAT expression levels were analyzed by total DAT/GAPDH ratio. (mean ± SEM, two-way ANOVA followed by Tukey test, * indicates p < 0.05; n = 3-5).

Even though total DAT expression levels remained invariable in both genotypes, DAT activity in the STR of p35KO mice was significantly lower than in WT controls, suggesting that the lack of Cdk5 activity exhibited by p35KO mice might affect transporters density at the plasma membrane.

### Mutant p35KO mice exhibit reduced DAT cell surface expression in the striatum

Taking into account the decreased DA reuptake observed in STR of p35KO animals, we investigate whether the lack of p35-activated Cdk5 activity exhibited by p35KO mice affects DAT surface expression. For this purpose, cell surface biotinylation experiments were performed on striatal synaptosomes obtained from WT and p35KO mice. Quantitative analysis of total extracts and surface fractions (biotinylated DAT/total DAT ratios) showed a 60 % reduction in the amount of surface DAT in p35KO mice compared to WT (Fig. 1c) (t-test: t = -3.007, df = 5, p < 0.05). As shown above, total DAT levels in the synaptosomal fraction remained invariable between genotypes (t-test: t = 2.180, df = 5, p = 0.081) (Fig. 1c). It is important to note the lack of GAPDH in biotinylated fractions ensuring they are free from intracellular contaminants (Fig. 1c).

We have previously shown that AMPH treatment induces a paradoxical calming effect on locomotor activity in p35KO mice and restores STR DA content to WT levels (Krapacher et al., 2010). Since AMPH exposure triggers DAT internalization (Johnson et al., 2005), we next analyzed the effect of this psychostimulant on DAT surface expression levels in striatal synaptosomes from p35KO and WT mice. Quantitative analysis revealed that 30 min of AMPH (10 µM) exposure significantly decreased DAT surface levels in synaptosomes from WT animals (two-way ANOVA, *F* _(1,11)_ = 17.684 *p* < 0.01) (Fig. 1d). However, incubation of synaptosomes from p35KO mice with AMPH did not affect DAT surface expression levels (p35KO vehicle compared to p35KO AMPH *p* = 0.490). Notably, vehicle incubated p35KO striatal synaptosomes displayed surface DAT expression levels similar to those observed in AMPH stimulated WT synaptosomes (WT AMPH vs p35KO Veh, *p* = 0.171). Finally, no significant statistical differences were observed in total DAT levels in synaptosomal fractions from both genotypes, regardless of the treatment given (two-way ANOVA, *F* _(1,11)_ = 2.651, *p=* 0.131) (Fig. 1d).

Taken together, our results demonstrate that mutant mice lacking p35, exhibit low DA uptake along with diminished surface DAT expression levels.

### Inhibition of Cdk5 increases DAT endocytosis in N2a cells

The dynamic balance between endocytosis and recycling is known to determine DAT surface levels. To determine whether Cdk5 participates in DAT internalization, we used the antibody-feeding method (AFM) in DAT-HA transfected N2a cells, under pharmacological Cdk5 activity inhibition with the specific inhibitor roscovitine (ROSCO). Since DAT endocytosis triggered by AMPH had been reported in a broad variety of cell lines, we used this psychostimulant as a positive control of DAT internalization (Chi and Reith, 2003; Saunders et al., 2000; Sorkina et al., 2003). The results showed that ROSCO (10 µM) or AMPH (10 µM) incubations in DAT-HA transfected N2a cells, significantly raised intracellular/surface DAT-HA ratios, demonstrating that inhibition of Cdk5 activity or AMPH treatment increased DAT endocytosis (Kruskal-Wallis test: H = 8.76, df = 2, p < 0.05) (Fig. 2a and b). Consistent with these observations, the expression of shRNA-mediated Cdk5 knockdown also increased intracellular/surface DAT-HA ratios (data not shown). These results suggest that Cdk5 might be involved in DAT internalization and provide more evidences to support functional and biochemical results obtained in p35KO mice.

**Figure 2.**
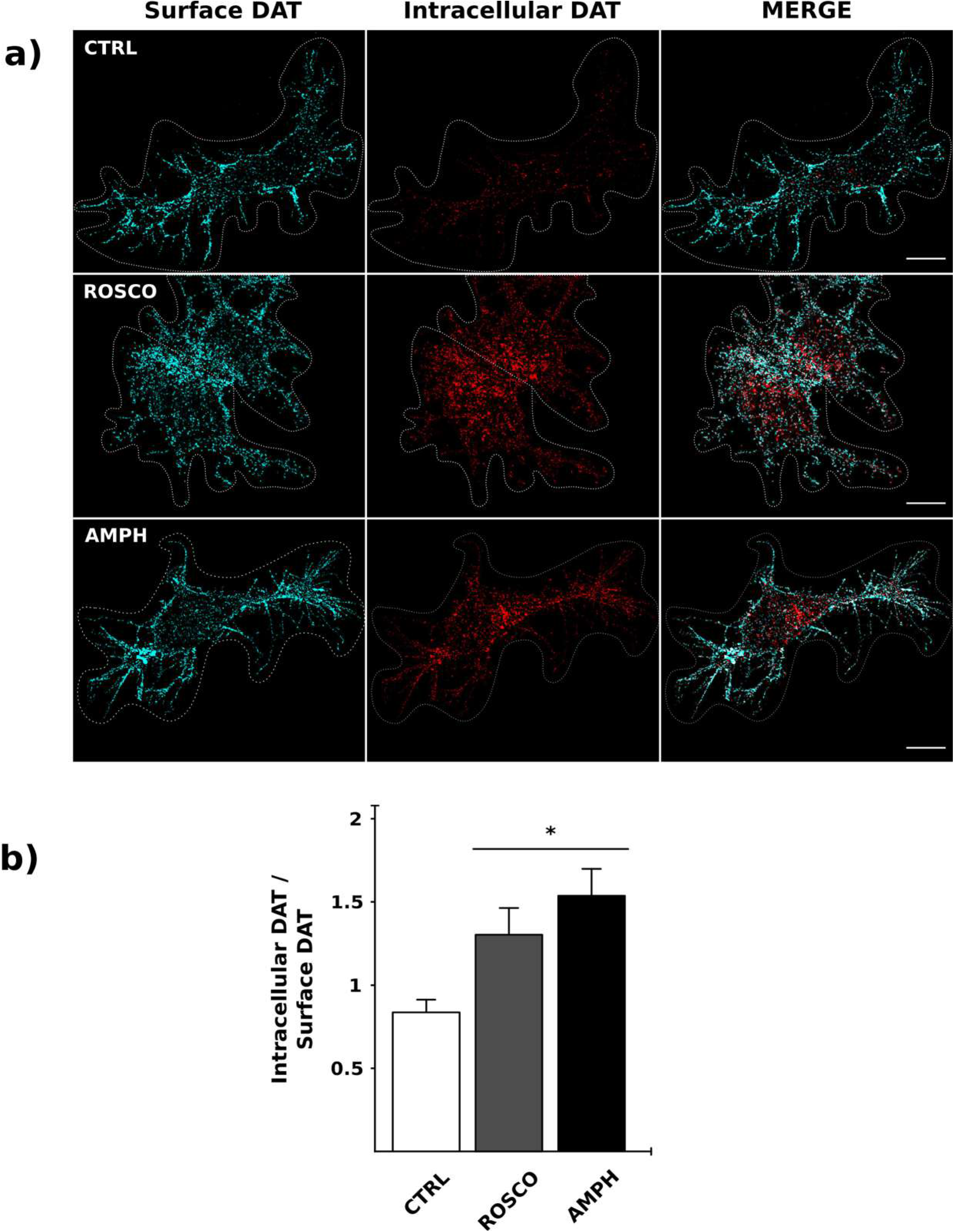
Inhibition of Cdk5 activity or AMPH treatment increases DAT internalization in N2a cells. (a) Cell surface DAT-HA was labeled by anti-HA11 followed by drug-induced internalization. Representative images of control (CTRL), ROSCO (10 µM) and AMPH (10 µM) treated N2a cells. Left panels show surface DAT-HA signal labeled with goat anti-rabbit-568, middle panels show internalized DAT-HA labeled with goat anti-rabbit-488 and right panels show the overlay of the two channels; scale bar = 10 µm. (b) Fluorescence quantification from confocal stacks between internalized DAT and surface DAT (mean ± SEM, Kruskal-Wallis test followed by Mann-Whitney U comparison tests; * indicates p < 0.05, n= 25-28 cells for each condition).

### Inhibition of Cdk5 activity arrests DAT in recycling endosomes

Taking into account that DAT endocytic pathway is highly regulated and involves several intermediate organelles, it is important to examine the fate of internalized DAT following Cdk5 activity inhibition and AMPH exposure in cultured N2a cells. For this purpose, we performed quantitative colocalization analysis of co-transfected N2a cells with DAT-HA with either Rab5-GFP or Rab11-GFP, early and recycling endosome markers, respectively (Stenmark, 2009). Using Manders’ Colocalization Coefficient (MCC) (Dunn et al., 2011), we observed a substantial constitutive colocalization (vehicle-control conditions) of intracellular DAT-HA with either Rab5-GFP or Rab11-GFP, which had similar MCC values (0.131 ± 0.019 and 0.154 ± 0.018, respectively) (Fig. 3a and b). Following 30 min of AMPH (10 µM) treatment, a significant amount of DAT-HA colocalized with both endosomal compartments (t-test: t = -3.491, df = 28, p < 0.01 for Rab5 positive endosomes and Kruskal-Wallis test: H = 10.961, df = 1, p < 0.01 for Rab11 positive endosomes). Noteworthy, AMPH incubation increased DAT-HA/Rab11-GFP MCC by approximately two-fold compared to DAT-HA/Rab5-GFP and almost three-fold compared to constitutively internalized DAT (Fig. 3b). These data are consistent with the enhanced DAT endocytosis observed in AMPH treated N2a cells (Fig. 2) and support the notion that internalized DAT following AMPH exposure traffics to early and recycling endosomes.

**Figure 3.**
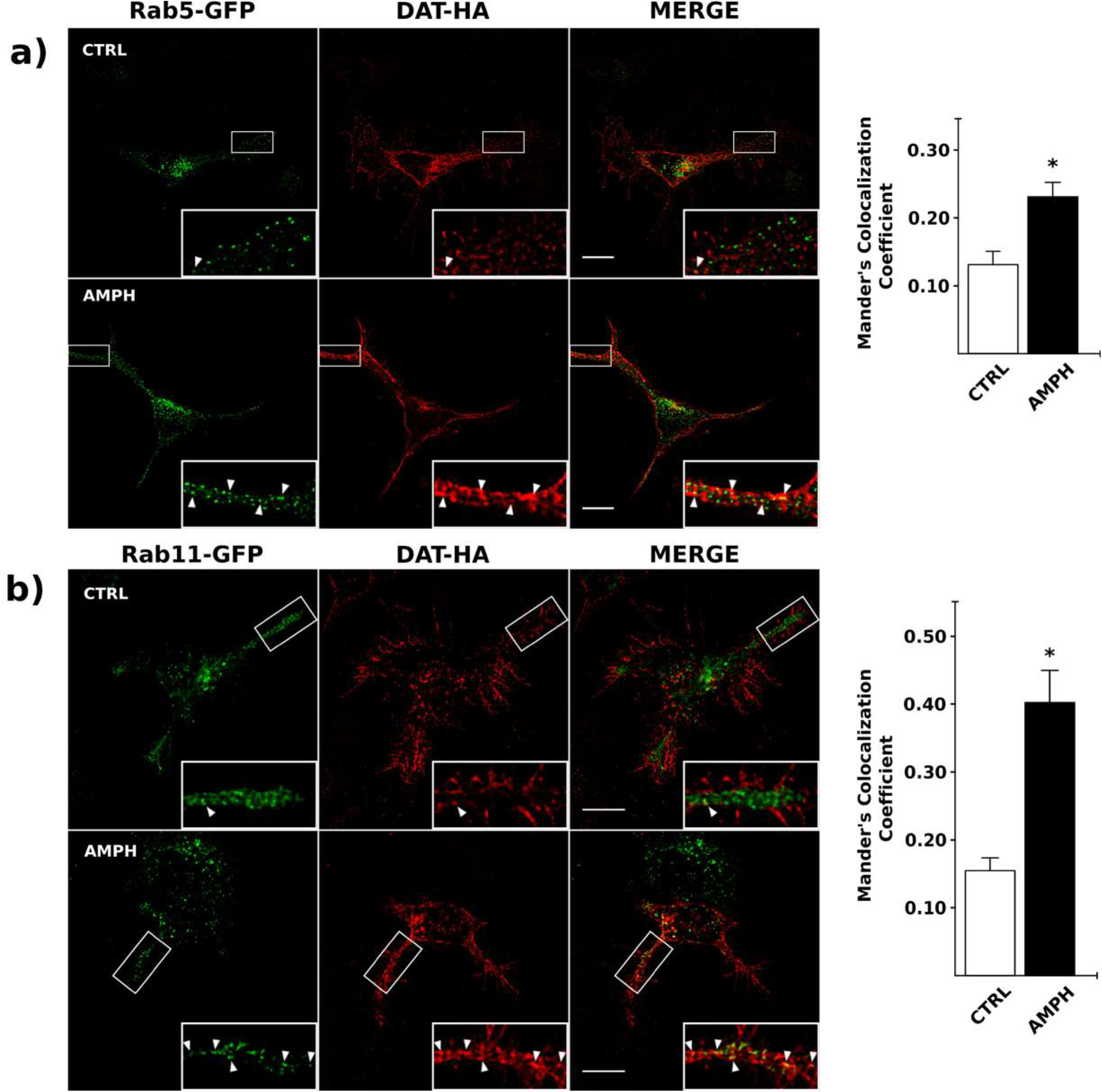
AMPH treatment increases DAT localization to early and recycling endosomes. N2a cells expressing DAT-HA and Rab5-GFP or Rab11-GFP were incubated with AMPH (10 µM) or vehicle (CTRL) for 30 min and colocalization between DAT and endosome markers was quantified. Representative Rab5-GFP/DAT-HA (a) and Rab11-GFP/DAT-HA (b) fluorescence signal images. Left panels show endosomal markers, medial panels show DAT-HA signal, and right panels, the overlay of the two channels. Bottom-right sections are enlarged images of boxed areas. Arrowheads indicate colocalized voxels. Scale bar = 10 µm. Bar graphs represent the quantification of fluorescence colocalization between DAT and endosome markers using Manders’ Colocalization Coefficient (MCC). (mean ± SEM, Student t-test in a and Kruskal-Wallis test in b, * indicates p < 0.05, n=10-16 processes for each condition).

Interestingly, when N2a cells co-transfected with DAT-HA and Rab5-GFP or Rab11-GFP were treated with ROSCO to inhibit Cdk5 activity, there was a dramatic change in DAT subcellular localization. In addition, a substantial amount of DAT-HA was observed only in recycling endosomes (Fig. 4). DAT-HA/Rab11-GFP MCC significantly increased compared to the vehicle-control condition, whereas DAT-HA/Rab5-GFP MCC remained invariable after the inhibition treatment (t-test: t = -2.869, df = 35, p < 0.01 for recycling Rab11 endosomes and t = -0.253, df = 46, p = 0.800 for early Rab5 endosomes) (Fig. 4a and b).

**Figure 4.**
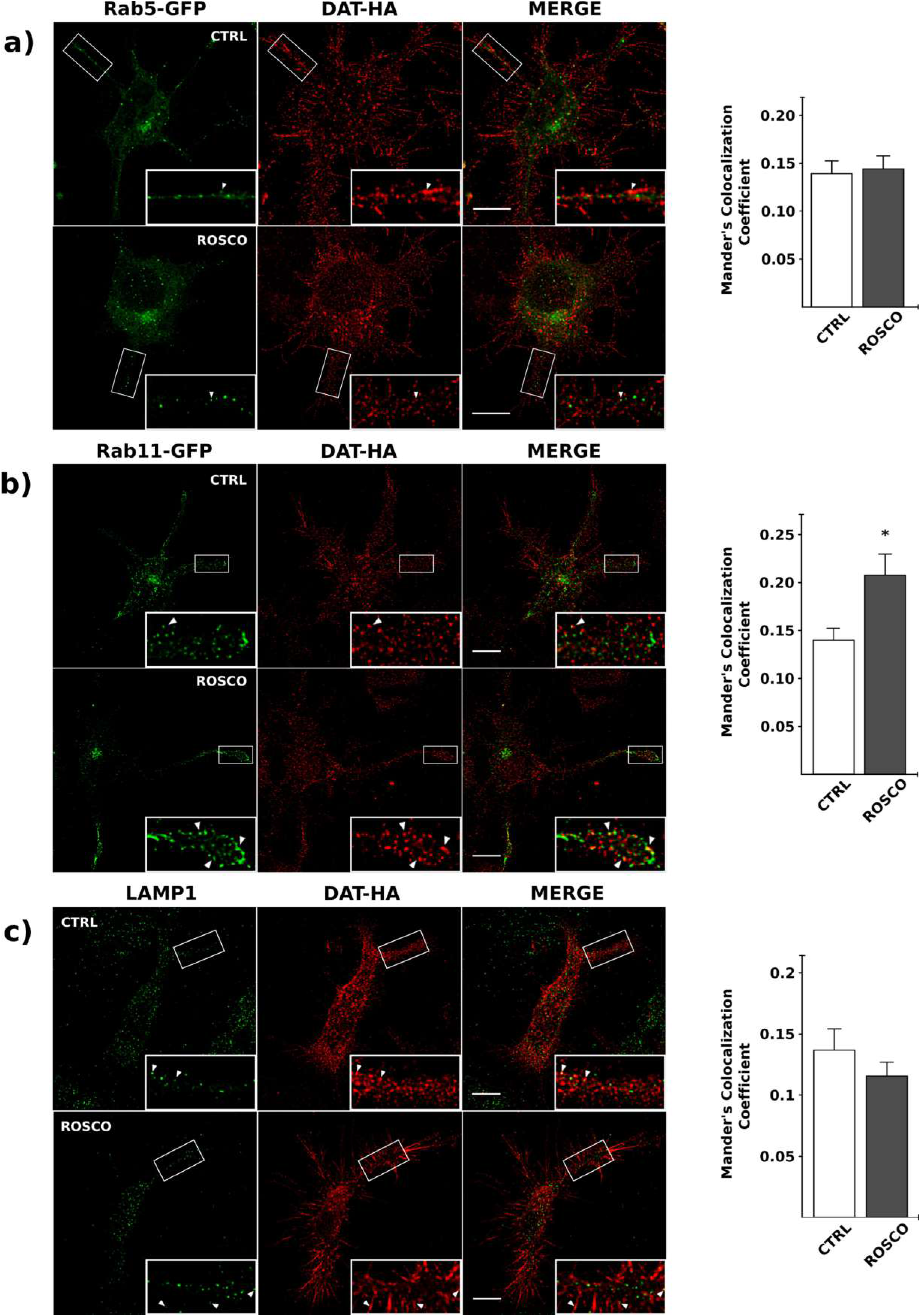
Cdk5 inhibition increases DAT localization only to recycling endosomes. N2a cells expressing DAT-HA or DAT-HA together with either Rab5-GFP or Rab11-GFP were treated with ROSCO (10 µM) or vehicle (CTRL) for 30 min. Then, colocalization between DAT with endosome and lysosome (LAMP-1) markers was quantified. Representative fluorescence signal images of Rab5-GFP/DAT-HA (a), Rab11-GFP/DAT-HA (b) and LAMP-1/DAT-HA (c). Left panels show endosomal and lysosomal markers, medial panels show DAT-HA signal, and right panels, the overlay of the two channels. Bottom-right sections are enlarged images of boxed areas. Arrowheads indicate colocalized voxels. Scale bar = 10 µm. Bar graphs represent quantification of fluorescence colocalization between DAT and endosome (a and b) and lysosome markers (c) using Manders’ Colocalization Coefficient (MCC) (mean ± SEM, Student t-test, * indicates p < 0.05 n=15-25 processes for each condition in a and b and n= 9-11 cells for each condition in c).

We next analyzed whether the fate of internalized DAT after Cdk5 activity inhibition is destined for degradation by analyzing colocalization of DAT-HA and LAMP1 (late endosome/lysosome marker) after 30 min of ROSCO exposure. According to the MCC quantification analysis, no significant differences were observed between DAT-HA/LAMP1 under Cdk5 inhibition and the vehicle-control condition (t-test: t = 1.120, df = 18, p = 0.277) (Fig. 4c).

These data suggest that pharmacological inhibition of Cdk5 activity increases constitutive DAT endocytosis, resulting in an accumulation of the transporter in recycling endosomes, maintaining the normal trafficking to the lysosomal degradative pathway.

## DISCUSSION

ADHD is one of the most common psychiatric conditions affecting both children and adults worldwide (Sayal et al., 2018). We have previously shown that the mutant mouse lacking p35 protein presents key hallmarks that resemble those of ADHD, thus supporting its validity as a genetically modified animal model useful for studying the disorder (de la Peña et al., 2017; Drerup et al., 2010; Krapacher et al., 2010). Besides, we also shown that p35KO presents increased TH protein levels, increased DA content and decreased DA degradation which result in a low neurotransmitter turnover (Krapacher et al., 2010). These features led us to hypothesize that the lack of p35-activated Cdk5 activity affects DAT expression and/or function in this animal model. In this study, we demonstrate that p35KO mice exhibit low DA uptake along with diminished surface DAT expression levels in the STR. These findings are supported by *in vitro* observations in which the inhibition of Cdk5 activity in N2a cells induced a significant increase in constitutive DAT endocytosis, with a concomitant loss of DAT from the cell surface and the increase of DAT localization to recycling endosomes.

The results obtained in p35KO mice show that the lack of p35-activated Cdk5 activity down-regulates striatal cell surface DAT without affecting the total expression levels of the transporter. Considering that pharmacological inhibition of Cdk5 kinase activity induces DAT internalization in N2a cells (Fig. 2), it is likely that an increase in DAT endocytic-rate underlies the reduced DAT surface levels observed in these mice. In addition, in previous studies we have demonstrated that inhibition of Cdk5 or p35 expression or its inactivation with ROSCO or olomoucine results in a decreased exocytic vesicles transport from the Golgi apparatus (Paglini et al., 2001), suggesting that the lack of Cdk5 activity may provoke an impairment of DAT trafficking to the cell surface. Furthermore, DAT losses from the plasma membrane are not due to blocking of newly synthesized DAT, because total DAT expression levels remained invariable between both genotypes. As expected, the reduction of striatal surface DAT in p35KO mice results in decreased DAT function. Our results are in agreement with previous reports on mutant mice expressing 10% of WT DAT levels (DAT knockdown) (Zhuang et al., 2001) and on PKCβ-knockout mice (Chen et al., 2009) as DA uptake in these mice is reduced. Decrease in DA uptake was also observed in rat striatal synaptosomes incubated *in vitro* with different Cdk5 inhibitors, in a concentration- and time-dependent manner (Price et al., 2009). Thus, our data suggest that p35-activated Cdk5 activity might be related to the maintenance of surface DAT in basal conditions, as evidenced by the reduced surface DAT expression level and DA uptake in STR of p35KO. Further experiments should be done to extend these observations to other brain areas implicated in the pathogenesis of ADHD, such as the prefrontal cortex. The spontaneous hyperactivity observed in p35KO mice (Krapacher et al., 2010) is consistent with the low surface DAT expression levels and reduced DA uptake reported here. Besides, our data are also in agreement with the increased basal locomotor activity described in DAT knockout and DAT knockdown mice (Gainetdinov, 1999; Zhuang et al., 2001).

The fact that AMPH belongs to the first line of ADHD pharmacological treatments (Briars and Todd, 2016), led a number of pharmacological studies to address the effect of this psychostimulant on DAT expression and activity. In this sense, AMPH treatment of striatal synaptosomes from WT mice reduced surface DAT expression levels. Likewise, the reduction in surface DAT upon exposure to AMPH has also been reported in rat and mouse STRs as well as in heterologous expression systems (Boudanova et al., 2008; Chen et al., 2009; Gulley et al., 2002; Johnson et al., 2005; Melikian, 2004; Saunders et al., 2000; Zahniser and Sorkin, 2004). Incubation of striatal synaptosomes from p35KO mice with AMPH elicits a decrease in surface DAT expression compared to vehicle. Even though this decrease was not statistically significant, it suggests that the lack of AMPH effect could reflect a floor effect, where surface DAT levels could not be reduced further or where the effect of AMPH is dependent on DAT superficial levels. Interestingly, vehicle-incubated p35KO striatal synaptosomes display low surface DAT expression levels similar to those observed in AMPH stimulated WT synaptosomes. The calming effect of AMPH on p35KO mice and the fact that synaptosomes of these mice present decreased DAT surface expression, are in line with previous studies indicating that AMPH behavioral effect on DAT transgenic mice is dependent on the degree of DAT expression (Cagniard et al., 2014; Salahpour et al., 2008). Previous studies have reported that the serotoninergic system is involved in the psychostimulant calming effect observed in ADHD animal models. For instance, methylendioximethamphetamine (MDMA), which has a higher effect on serotoninergic relative to dopaminergic neurons, decreases locomotor activity in DAT-knockdown mice (Cagniard et al., 2014). Moreover, the serotonin inhibitor fluoxetine exerts a calming effect on DAT-KO mice (Gainetdinov, 1999). Therefore, given the low DAT surface expression in p35KO mice, the serotoninergic system is likely to be involved in the AMPH paradoxical effect observed in these animals. Future research should focus on determining the role of serotoninergic system on the behavioral phenotype as well as the neurochemical patterns in p35KO mice.

Performing *in vitro* experiments, using N2a cells, we demonstrated that pharmacological inhibition of Cdk5 kinase activity with the widely used inhibitor ROSCO increases DAT internalization (Fig. 2), suggesting that Cdk5 could be acting as a negative regulator of constitutive DAT endocytosis, supporting functional and biochemical results obtained in p35KO mice. It is well known that the amount of cell surface DAT is the result of a dynamic balance between endocytosis and recycling to the membrane, processes that are highly regulated by a variety of proteins. In this sense, it has been demonstrated that constitutive and substrate induced DAT internalization are clathrin-dependent processes, where dynamin is involved (Carvelli et al., 2002; Damke et al., 1994; Saunders et al., 2000; Sorkina et al., 2005). Interestingly, Cdk5-mediated phosphorylation of dynamin I impairs its association with amphiphysin I, resulting in synaptic vesicle endocytosis inhibition. Moreover, evidences obtained in p35KO mice support these findings since vesicle endocytosis is enhanced in hippocampal neurons from these animals (Tomizawa et al., 2003). Our *in vitro* and *in vivo* results reinforce and extend these observations since p35KO mice exhibit a reduced cell surface DAT expression in the STR.

Because internalized DAT-containing vesicles have a specific postendocytic sorting (Eriksen et al., 2010; Hong and Amara, 2013), we analyzed the internalization pathways of the transporter under Cdk5 inhibition or AMPH treatment in N2a cells. Under steady-state conditions, we observed a substantial colocalization of DAT with both, early (Rab5 positive) and recycling (Rab11 positive) endosomes. These results are consistent with previous reports demonstrating that internalized DAT is sorted to the recycling pathways (Loder and Melikian, 2003; Sorkina et al., 2005). In addition, AMPH incubation not only increases DAT endocytosis, but also significantly increases DAT localization to early and recycling endosomes (Fig. 3). These results support the notion that internalized DAT, under stimulant conditions, traffics to the recycling endosome pathway for surface reinsertion, as previously shown in cultured MN9D and HEK293 cell lines and in embryonic rat primary mesencephalic cultures (Hong and Amara, 2013). We next examined the fate of internalized DAT following pharmacological inhibition of Cdk5 kinase activity in cultured N2a cells. Surprisingly, ROSCO treatment enhances only DAT-recycling endosomes colocalization. This might indicates that the lack of Cdk5 activity induces the sorting of constitutive internalized-DAT to the recycling pathway. Some lines of evidence support this notion. Small GTPase Rab11 and its effectors control recycling endosomes trafficking and also transporters and receptors replenishment at the plasma membrane (Johnson et al., 2016). Moreover, a recent report showed that Cdk5/p35 complex regulates the Rab11-effector GRAB by phosphorylation, modulating the transport of Rab11-dependent endosomes (Furusawa et al., 2017). Furthermore, the final step of the trafficking process is the docking and fusion of Rab11-positive vesicles at the plasma membrane, where the involvement of Cdk5 has already been demonstrated (Liu et al., 2007; Tomizawa et al., 2002).

Taken all together these data provide new evidences of dopaminergic neurotransmission alterations in p35KO mice and suggest the involvement of Cdk5 in DAT cell surface availability. The high ADHD prevalence has raised the interest of the scientific community to understanding its neurobiological bases. In this context, animal models of human diseases play a critical role in the exploration and characterization of the pathophysiology and represent a very useful tool in preclinical research to identify, evaluate and validate novel therapeutic agents and treatments.

## CONCLUSIONS

Overall, this study provides new insights regarding the role of Cdk5/p35 in the maintenance of DAT at the plasma membrane for its functional availability. In addition, this research extends the knowledge of p35KO mouse dopaminergic system and its modulation to provide a better understanding of the molecular mechanisms underlying ADHD. Nowadays, it is known that ADHD is a chronic condition that continues into adulthood and psychostimulant treatments are often prescribed for years. Therefore, the availability of animal models combined with in vitro approaches are invaluable tools to unscramble the complicated nature of complex psychiatric disorders like ADHD, and provide a great opportunity to screen potential therapeutic strategies.

## ABBREVIATIONS

ADHD: Attention deficit/Hyperactivity disorder
AFM: antibody feeding method
AMPH: amphetamine
DA: Dopamine
DAT-HA: DAT containing a HA epitope in the second extracellular loop
DAT: Dopamine Transporter
GAPDH: glyceraldehyde 3-phosphate dehydrogenase
GRAB: guanine nucleotide exchange factor for Rab
LAMP-1: Lysosome-associated membrane protein 1
MCC: Manders’ Colocalization Coefficient
p35KO: mice lacking p35 protein
ROSCO: roscovitine
STR: striatum
WT: wild type mice

## Authors’ contributions

All authors were responsible for the design of the study. GF, FK and SF performed the experimental procedures and collected data. MVP and CA conducted statistical data analysis. GF and GQ performed imaging analysis. GF and MMM conducted biochemical essays. AM and MDR performed amperometric experiments. CB and CA contributed ideas and comments. GF and MGP interpreted the findings and drafted the manuscript. Supervision and project administration were done by MGP and funding’s acquisition by MGP and CA. All authors critically reviewed contents and approved the final version for publication.

## Acknowledgements

We thank Maria Julia Cambiasso for her all-encompassing support, Andrea Pellegrini for technical assistance, Mariana Bollo, Pablo Helguera and Mariano Bisbal for generously providing some antibodies. We would particularly like to thank Damian Revillo and Evelin Cotella for helpful discussions, Patricio Pereyra for his assistance with animal husbandry and the vivarium technicians for their daily support. Confocal microscopy was performed at the Centro de Micro y Nanoscopía de Córdoba, CEMINCO-CONICET-Universidad Nacional de Córdoba, Córdoba, Argentina.

## REFERENCES

American Psychiatric Association, 2013. DSM-5 Diagnostic Classification, in: Diagnostic and Statistical Manual of Mental Disorders. American Psychiatric Association. https://doi.org/10.1176/appi.books.9780890425596.x00DiagnosticClassification

Bermingham, D.P., Blakely, R.D., 2016. Kinase-dependent Regulation of Monoamine Neurotransmitter Transporters. Pharmacol. Rev. 68, 888–953. https://doi.org/10.1124/pr.115.012260

Bolte, S., Cordelières, F.P., 2006. A guided tour into subcellular colocalization analysis in light microscopy. J. Microsc. 224, 213–32. https://doi.org/10.1111/j.1365-2818.2006.01706.x

Boudanova, E., Navaroli, D.M., Melikian, H.E., 2008. Amphetamine-induced decreases in dopamine transporter surface expression are protein kinase C-independent. Neuropharmacology 54, 605–612. https://doi.org/10.1016/j.neuropharm.2007.11.007

Briars, L., Todd, T., 2016. A Review of Pharmacological Management of Attention-Deficit/Hyperactivity Disorder. J. Pediatr. Pharmacol. Ther. 21, 192–206. https://doi.org/10.5863/1551-6776-21.3.192

Cagniard, B., Sotnikova, T.D., Gainetdinov, R.R., Zhuang, X., 2014. The Dopamine Transporter Expression Level Differentially Affects Responses to Cocaine and Amphetamine. J. Neurogenet. 28, 112–121. https://doi.org/10.3109/01677063.2014.908191

Carvelli, L., Morón, J.A., Kahlig, K.M., Ferrer, J. V., Sen, N., Lechleiter, J.D., Leeb-Lundberg, L.M.F., Merrill, G., Lafer, E.M., Ballou, L.M., Shippenberg, T.S., Javitch, J.A., Lin, R.Z., Galli, A., 2002. PI 3-kinase regulation of dopamine uptake. J. Neurochem. 81, 859–869. https://doi.org/10.1046/j.1471-4159.2002.00892.x

Chae, T., Kwon, Y.T., Bronson, R., Dikkes, P., Li, E., Tsai, L.-H., 1997. Mice Lacking p35, a Neuronal Specific Activator of Cdk5, Display Cortical Lamination Defects, Seizures, and Adult Lethality. Neuron 18, 29–42. https://doi.org/10.1016/S0896-6273(01)80044-1

Chen, R., Furman, C.A., Zhang, M., Kim, M.N., Gereau, R.W., Leitges, M., Gnegy, M.E., 2009. Protein Kinase Cβ Is a Critical Regulator of Dopamine Transporter Trafficking and Regulates the Behavioral Response to Amphetamine in Mice. J. Pharmacol. Exp. Ther. 328, 912–920. https://doi.org/10.1124/jpet.108.147959

Chi, L., Reith, M.E. a, 2003. Substrate-induced trafficking of the dopamine transporter in heterologously expressing cells and in rat striatal synaptosomal preparations. J. Pharmacol. Exp. Ther. 307, 729–736. https://doi.org/10.1124/jpet.103.055095

Damke, H., Baba, T., Warnock, D.E., Schmid, S.L., 1994. Induction of mutant dynamin specifically blocks endocytic coated vesicle formation. J. Cell Biol. 127, 915–934. https://doi.org/10.1083/jcb.127.4.915

de la Peña, J.B., dela Peña, I.J., Custodio, R.J., Botanas, C.J., Kim, H.J., Cheong, J.H., 2017. Exploring the Validity of Proposed Transgenic Animal Models of Attention-Deficit Hyperactivity Disorder (ADHD). Mol. Neurobiol. 55, 3739–3754. https://doi.org/10.1007/s12035-017-0608-1

Drerup, J.M., Hayashi, K., Cui, H., Mettlach, G.L., Long, M.A., Marvin, M., Sun, X., Goldberg, M.S., Lutter, M., Bibb, J.A., 2010. Attention-deficit/hyperactivity phenotype in mice lacking the cyclin-dependent kinase 5 cofactor p35. Biol. Psychiatry 68, 1163–1171. https://doi.org/10.1016/j.biopsych.2010.07.016

Dunn, K.W., Kamocka, M.M., McDonald, J.H., 2011. A practical guide to evaluating colocalization in biological microscopy. AJP Cell Physiol. 300, C723–C742. https://doi.org/10.1152/ajpcell.00462.2010

Eriksen, J., Bjørn-Yoshimoto, W.E., Jørgensen, T.N., Newman, A.H., Gether, U., 2010. Postendocytic sorting of constitutively internalized dopamine transporter in cell lines and dopaminergic neurons. J. Biol. Chem. 285, 27289–27301. https://doi.org/10.1074/jbc.M110.131003

Faraone, S. V., Perlis, R.H., Doyle, A.E., Smoller, J.W., Goralnick, J.J., Holmgren, M.A., Sklar, P., 2005. Molecular genetics of attention-deficit/hyperactivity disorder. Biol. Psychiatry 57, 1313–1323. https://doi.org/10.1016/j.biopsych.2004.11.024

Federici, M., Latagliata, E.C., Ledonne, A., Rizzo, F.R., Feligioni, M., Sulzer, D., Dunn, M., Sames, D., Gu, H., Nisticò, R., Puglisi-Allegra, S., Mercuri, N.B., 2014. Paradoxical Abatement of Striatal Dopaminergic Transmission by Cocaine and Methylphenidate. J. Biol. Chem. 289, 264–274. https://doi.org/10.1074/jbc.M113.495499

Ferreras, S., Fernández, G., Danelon, V., Pisano, M. V., Masseroni, L., Chapleau, C.A., Krapacher, F.A., Mlewski, E.C., Mascó, D.H., Arias, C., Pozzo-Miller, L., Paglini, M.G., 2017. Cdk5 Is Essential for Amphetamine to Increase Dendritic Spine Density in Hippocampal Pyramidal Neurons. Front. Cell. Neurosci. 11. https://doi.org/10.3389/fncel.2017.00372

Furusawa, K., Asada, A., Urrutia, P., Gonzalez-Billault, C., Fukuda, M., Hisanaga, S., 2017. Cdk5 Regulation of the GRAB-Mediated Rab8-Rab11 Cascade in Axon Outgrowth. J. Neurosci. 37, 790–806. https://doi.org/10.1523/JNEUROSCI.2197-16.2016

Gainetdinov, R.R., 1999. Role of Serotonin in the Paradoxical Calming Effect of Psychostimulants on Hyperactivity. Science (80-.). 283, 397–401. https://doi.org/10.1126/science.283.5400.397

German, C.L., Baladi, M.G., McFadden, L.M., Hanson, G.R., Fleckenstein, A.E., 2015. Regulation of the Dopamine and Vesicular Monoamine Transporters: Pharmacological Targets and Implications for Disease. Pharmacol. Rev. 67, 1005–1024. https://doi.org/10.1124/pr.114.010397

Gulley, J.M., Doolen, S., Zahniser, N.R., 2002. Brief, repeated exposure to substrates down-regulates dopamine transporter function in Xenopus oocytes in vitro and rat dorsal striatum in vivo. J. Neurochem. 83, 400–411. https://doi.org/10.1046/j.1471-4159.2002.01133.x

Hawi, Z., Cummins, T.D.R., Tong, J., Johnson, B., Lau, R., Samarrai, W., Bellgrove, M.A., 2015. The molecular genetic architecture of attention deficit hyperactivity disorder. Mol. Psychiatry 20, 289–297. https://doi.org/10.1038/mp.2014.183

Hong, W.C., Amara, S.G., 2013. Differential targeting of the dopamine transporter to recycling or degradative pathways during amphetamine-or PKC-regulated endocytosis in dopamine neurons. FASEB J. 27, 2995–3007. https://doi.org/10.1096/fj.12-218727

Ikiz, B., Przedborski, S., 2008. Previews A Sequel to the Tale of p25 / Cdk5 in Neurodegeneration 731–732. https://doi.org/10.1016/j.neuron.2008.11.020

Ishiguro, K., Kobayashi, S., Onion, A., Takamatsu, M., Yonekura, S., Anzai, K., Imahori, K., Uchida, T., 1994. Identification of the 23 kDa subunit of tau protein kinase II as a putative activator of cdk5 in bovine brain. FEBS Lett. https://doi.org/10.1016/0014-5793(94)80501-6

Johnson, J.L., He, J., Ramadass, M., Pestonjamasp, K., Kiosses, W.B., Zhang, J., Catz, S.D., 2016. Munc13-4 Is a Rab11-binding Protein That Regulates Rab11-positive Vesicle Trafficking and Docking at the Plasma Membrane. J. Biol. Chem. 291, 3423–3438. https://doi.org/10.1074/jbc.M115.705871

Johnson, L.A., Furman, C.A., Zhang, M., Guptaroy, B., Gnegy, M.E., 2005. Rapid delivery of the dopamine transporter to the plasmalemmal membrane upon amphetamine stimulation. Neuropharmacology 49, 750–758. https://doi.org/10.1016/j.neuropharm.2005.08.018

Kawauchi, T., 2014. Cdk5 regulates multiple cellular events in neural development, function and disease. Dev. Growth Differ. 56, 335–348. https://doi.org/10.1111/dgd.12138

Kim, S.H., Ryan, T.A., 2010. CDK5 Serves as a Major Control Point in Neurotransmitter Release. Neuron 67, 797–809. https://doi.org/10.1016/j.neuron.2010.08.003

Krapacher, F.A., Mlewski, E.C., Ferreras, S., Pisano, V., Paolorossi, M., Hansen, C., Paglini, G., 2010. Mice lacking p35 display hyperactivity and paradoxical response to psychostimulants. J. Neurochem. 114, 203–14. https://doi.org/10.1111/j.1471-4159.2010.06748.x

Leo, D., Gainetdinov, R.R., 2013. Transgenic mouse models for ADHD. Cell Tissue Res. 354, 259–271. https://doi.org/10.1007/s00441-013-1639-1

Lew, J., Wang, J.H., 1995. Neuronal cdc2-like kinase. Trends Biochem. Sci. https://doi.org/10.1016/S0968-0004(00)88948-3

Li, Z., Chang, S. hua Zhang, L. yan, Gao, L., Wang, J., 2014. Molecular genetic studies of ADHD and its candidate genes: A review. Psychiatry Res. https://doi.org/10.1016/j.psychres.2014.05.005

Liu, Y., Ding, X., Wang, D., Deng, H., Feng, M., Wang, M., Yu, X., Jiang, K., Ward, T., Aikhionbare, F., Guo, Z., Forte, J.G., Yao, X., 2007. A mechanism of Munc18b-syntaxin 3-SANP25 complex assembly in regulated epithelial secretion. FEBS Lett. 581, 4318–4324. https://doi.org/10.1016/j.febslet.2007.07.083

Loder, M.K., Melikian, H.E., 2003. The dopamine transporter constitutively internalizes and recycles in a protein kinase C-regulated manner in stably transfected PC12 cell lines. J. Biol. Chem. 278, 22168–74. https://doi.org/10.1074/jbc.M301845200

López-Maderuelo, M.D., Fernández-Renart, M., Moratilla, C., Renart, J., 2001. Opposite effects of the Hsp90 inhibitor Geldanamycin: induction of apoptosis in PC12, and differentiation in N2A cells. FEBS Lett. 490, 23–7. https://doi.org/10.1016/s0014-5793(01)02130-5

Melikian, H.E., 2004. Neurotransmitter transporter trafficking: endocytosis, recycling, and regulation. Pharmacol. Ther. 104, 17–27. https://doi.org/10.1016/j.pharmthera.2004.07.006

Mlewski, E.C., Arias, C., Paglini, G., 2016. Association between the expression of amphetamine-induced behavioral sensitization and Cdk5/p35 activity in dorsal striatum. Behav. Neurosci. 130. https://doi.org/10.1037/bne0000118

Mlewski, E.C., Krapacher, F.A., Ferreras, S., Paglini, G., 2008. Transient Enhanced Expression of Cdk5 Activator p25 after Acute and Chronic d -Amphetamine Administration. Ann. N. Y. Acad. Sci. 1139, 89–102. https://doi.org/10.1196/annals.1432.039

Napolitano, F., Bonito-Oliva, A., Federici, M., Carta, M., Errico, F., Magara, S., Martella, G., Nisticò, R., Centonze, D., Pisani, A., Gu, H.H., Mercuri, N.B., Usiello, A., 2010. Role of aberrant striatal dopamine D1 receptor/cAMP/protein kinase A/DARPP32 signaling in the paradoxical calming effect of amphetamine. J. Neurosci. 30, 11043–56. https://doi.org/10.1523/JNEUROSCI.1682-10.2010

Nguyen, C., Bibb, J.A., 2003. Cdk5 and the mystery of synaptic vesicle endocytosis. J. Cell Biol. 163, 697–699. https://doi.org/10.1083/jcb.200310038

O’Neill, B., Gu, H.H., 2013. Amphetamine-induced locomotion in a hyperdopaminergic ADHD mouse model depends on genetic background. Pharmacol. Biochem. Behav. 103, 455–459. https://doi.org/10.1016/j.pbb.2012.09.020

Paglini, G., Peris, L., Diez-Guerra, J., Quiroga, S., Cáceres, A., 2001. The Cdk5-p35 kinase associates with the Golgi apparatus and regulates membrane traffic. EMBO Rep. 2, 1139–44. https://doi.org/10.1093/embo-reports/kve250

Paglini, G., Pigino, G., Kunda, P., Morfini, G., Maccioni, R., Quiroga, S., Ferreira, a, Cáceres, a, 1998. Evidence for the participation of the neuron-specific CDK5 activator P35 during laminin-enhanced axonal growth. J. Neurosci. 18, 9858–9869.

Piccini, A., Perlini, L.E., Cancedda, L., Benfenati, F., Giovedi, S., 2015. Phosphorylation by PKA and Cdk5 Mediates the Early Effects of Synapsin III in Neuronal Morphological Maturation. J. Neurosci. 35, 13148–13159. https://doi.org/10.1523/JNEUROSCI.1379-15.2015

Price, D.A., Sorkin, A., Zahniser, N.R., 2009. Cyclin-dependent kinase 5 inhibitors: inhibition of dopamine transporter activity. Mol Pharmacol 76, 812–823. https://doi.org/10.1124/mol.109.056978

R Core Team, 2020. R: A language and environment for statistical computing.

Rei, D., Mason, X., Seo, J., Gräff, J., Rudenko, A., Wang, J., Rueda, R., Siegert, S., Cho, S., Canter, R.G., Mungenast, A.E., Deisseroth, K., Tsai, L.H., 2015. Basolateral amygdala bidirectionally modulates stress-induced hippocampal learning and memory deficits through a p25/Cdk5-dependent pathway. Proc. Natl. Acad. Sci. U. S. A. https://doi.org/10.1073/pnas.1415845112

Rubianes, M.D., Rivas, G.A., 2007. Dispersion of multi-wall carbon nanotubes in polyethylenimine: A new alternative for preparing electrochemical sensors. Electrochem. commun. 9, 480–484. https://doi.org/10.1016/j.elecom.2006.08.057

Sage, D., Donati, L., Soulez, F., Fortun, D., Schmit, G., Seitz, A., Guiet, R., Vonesch, C., Unser, M., 2017. DeconvolutionLab2: An open-source software for deconvolution microscopy. Methods 115, 28–41. https://doi.org/10.1016/j.ymeth.2016.12.015

Sakrikar, D., Mazei-Robison, M.S., Mergy, M.A., Richtand, N.W., Han, Q., Hamilton, P.J., Bowton, E., Galli, A., Veenstra-VanderWeele, J., Gill, M., Blakely, R.D., 2012. Attention Deficit/Hyperactivity Disorder-Derived Coding Variation in the Dopamine Transporter Disrupts Microdomain Targeting and Trafficking Regulation. J. Neurosci. 32, 5385–5397. https://doi.org/10.1523/JNEUROSCI.6033-11.2012

Salahpour, A., Ramsey, A.J., Medvedev, I.O., Kile, B., Sotnikova, T.D., Holmstrand, E., Ghisi, V., Nicholls, P.J., Wong, L., Murphy, K., Sesack, S.R., Wightman, R.M., Gainetdinov, R.R., Caron, M.G., 2008. Increased amphetamine-induced hyperactivity and reward in mice overexpressing the dopamine transporter. Proc. Natl. Acad. Sci. U. S. A. 105, 4405–10. https://doi.org/10.1073/pnas.0707646105

Saunders, C., Ferrer, J. V., Shi, L., Chen, J., Merrill, G., Lamb, M.E., Leeb-Lundberg, L.M.F., Carvelli, L., Javitch, J.A., Galli, A., 2000. Amphetamine-induced loss of human dopamine transporter activity: An internalization-dependent and cocaine-sensitive mechanism. Proc. Natl. Acad. Sci. 97, 6850–6855. https://doi.org/10.1073/pnas.110035297

Sayal, K., Prasad, V., Daley, D., Ford, T., Coghill, D., 2018. ADHD in children and young people: prevalence, care pathways, and service provision. The Lancet Psychiatry. https://doi.org/10.1016/S2215-0366(17)30167-0

Shah, K., Lahiri, D.K., 2017. A Tale of the Good and Bad: Remodeling of the Microtubule Network in the Brain by Cdk5. Mol. Neurobiol. 54, 2255–2268. https://doi.org/10.1007/s12035-016-9792-7

Shah, K., Lahiri, D.K., 2014. Cdk5 activity in the brain-multiple paths of regulation. J. Cell Sci. 127, 2391–400. https://doi.org/10.1242/jcs.147553

Shah, K., Rossie, S., 2018. Tale of the Good and the Bad Cdk5: Remodeling of the Actin Cytoskeleton in the Brain. Mol. Neurobiol. 55, 3426–3438. https://doi.org/10.1007/s12035-017-0525-3

Sharma, A., Couture, J., 2014. A Review of the Pathophysiology, Etiology, and Treatment of Attention-Deficit Hyperactivity Disorder (ADHD). Ann. Pharmacother. 48, 209–225. https://doi.org/10.1177/1060028013510699

Sorkina, T., Doolen, S., Galperin, E., Zahniser, N.R., Sorkin, A., 2003. Oligomerization of dopamine transporters visualized in living cells by fluorescence resonance energy transfer microscopy. J. Biol. Chem. 278, 28274–28283. https://doi.org/10.1074/jbc.M210652200

Sorkina, T., Hoover, B.R., Zahniser, N.R., Sorkin, A., 2005. Constitutive and Protein Kinase C-Induced Internalization of the Dopamine Transporter is Mediated by a Clathrin-Dependent Mechanism. Traffic 6, 157–170. https://doi.org/10.1111/j.1600-0854.2005.00259.x

Sorkina, T., Miranda, M., Dionne, K.R., Hoover, B.R., Zahniser, N.R., Sorkin, A., 2006. RNA interference screen reveals an essential role of Nedd4-2 in dopamine transporter ubiquitination and endocytosis. J. Neurosci. 26, 8195–8205. https://doi.org/10.1523/JNEUROSCI.1301-06.2006

Stenmark, H., 2009. Rab GTPases as coordinators of vesicle traffic. Nat. Rev. Mol. Cell Biol. 10, 513–25. https://doi.org/10.1038/nrm2728

Svenningsson, P., Nishi, A., Fisone, G., Girault, J.-A., Nairn, A.C., Greengard, P., 2004. DARPP-32: An Integrator of Neurotransmission. Annu. Rev. Pharmacol. Toxicol. 44, 269–296. https://doi.org/10.1146/annurev.pharmtox.44.101802.121415

Tan, T.C., Valova, V. a, Malladi, C.S., Graham, M.E., Berven, L. a, Jupp, O.J., Hansra, G., McClure, S.J., Sarcevic, B., Boadle, R. a, Larsen, M.R., Cousin, M. a, Robinson, P.J., 2003. Cdk5 is essential for synaptic vesicle endocytosis. Nat. Cell Biol. 5, 701–710. https://doi.org/10.1038/ncb1020

Tang, D., Yeung, J., Lee, K.Y., Matsushita, M., Matsui, H., Tomizawa, K., Hatase, O., Wang, J.H., 1995. An isoform of the neuronal cyclin-dependent kinase 5 (Cdk5) activator. J. Biol. Chem. 270, 26897–26903. https://doi.org/10.1074/jbc.270.45.26897

Tomizawa, K., Ohta, J., Matsushita, M., Moriwaki, A., Li, S., Takei, K., Matsui, H., 2002. Cdk5/p35 regulates neurotransmitter release through phosphorylation and downregulation of P/Q-type voltage-dependent calcium channel activity. J. Neurosci. 22, 2590–7. https://doi.org/20026252

Tomizawa, K., Sunada, S., Lu, Y.F., Oda, Y., Kinuta, M., Ohshima, T., Saito, T., Wei, F.Y., Matsushita, M., Li, S.T., Tsutsui, K., Hisanaga, S.I., Mikoshiba, K., Takei, K., Matsui, H., 2003. Cophosphorylation of amphiphysin I and dynamin I by Cdk5 regulates clathrin-mediated endocytosis of synaptic vesicles. J. Cell Biol. 163, 813–824. https://doi.org/10.1083/jcb.200308110

Tsai, L.-H., Delalle, I., Caviness, V.S., Chae, T., Harlow, E., 1994. p35 is a neural-specific regulatory subunit of cyclin-dependent kinase 5. Nature 371, 419–423. https://doi.org/10.1038/371419a0

Zahniser, N.R., Sorkin, A., 2004. Rapid regulation of the dopamine transporter: Role in stimulant addiction? Neuropharmacology 47, 80–91. https://doi.org/10.1016/j.neuropharm.2004.07.010

Zhuang, X., Oosting, R.S., Jones, S.R., Gainetdinov, R.R., Miller, G.W., Caron, M.G., Hen, R., 2001. Hyperactivity and impaired response habituation in hyperdopaminergic mice. Proc. Natl. Acad. Sci. U. S. A. 98, 1982–7. https://doi.org/10.1073/pnas.98.4.1982

